# Critical dynamics arise during structured information presentation: analysis of embodied *in vitro* neuronal networks

**DOI:** 10.1101/2022.11.03.514955

**Authors:** Forough Habibollahi, Brett J. Kagan, Daniela Duc, Anthony N. Burkitt, Chris French

## Abstract

Amongst the characteristics about information processing in the brain, observations of dynamic near-critical states have generated significant interest. However, theoretical and experimental limitations have precluded a definite answer about when and why neural criticality arises. To explore this topic, we used an *in vitro* neural network of cortical neurons that was trained to play a simplified game of ‘Pong’. We demonstrate that critical dynamics emerge when neural networks receive task-related structured sensory input, reorganizing the system to a near-critical state. Additionally, better task performance correlated with proximity to critical dynamics. However, criticality alone is insufficient for a neuronal network to demonstrate learning in the absence of additional information regarding the consequences of previous actions. These findings have compelling implications for the role of neural criticality.

## Introduction

How do our brains process information? It has been hypothesised for some decades that neural systems operate in or near a “critical state” [1–6] with well-defined dynamical properties characterised by, inter alia, stability of neuronal activity, optimised information storage, and transmission [4, 7]. The presence of *neuronal avalanches* (cascades of propagating activity governed by power laws) that are one hallmark of criticality is widely reported in the spontaneous activity of *in vivo* cortical networks [8–12]. Evidence of neuronal avalanches have also been found *in vitro* in local field potentials (LFPs) of spontaneous activity in acute cortex slices [5, 13], slice cultures [5], cultured hippocampal neurons [14], and cultured human stem cells [15].Yet the specific role of neural criticality, along with why and when it occurs, is a matter of significant controversy [16].

Early work identified a link between the balance of excitatory and inhibitory inputs and the critical phase transition[17]. Anticipating this critical transition and the proximity of a network to criticality informs network robustness and can even approximate risk factors of network failures such as epileptic seizures [18]. Moreover, cortical networks express a dynamic equilibrium regime associated with criticality, including: 1) the absence of runaway gains, in which balanced activity is maintained in the neuronal networks such that the neuronal activity does not saturate or become quiescent [3]; 2) a wide coverage in both spatial (mm to cm) and temporal (ms to min, h, etc.) scales during information encoding and transmission [3]; 3) wide dynamical range [19, 20]; and 4) maximized information transmission in terms of mutual information [21, 22] and information storage and processing capabilities [23], such as elevated sensitivity and susceptibility to input. While in the context of population dynamics these criteria have been postulated to be a homeostatic set point for biological neural networks (BNNs), questions about the utility remain. [24–27].

Previous modelling of neuronal avalanches and of cortical slice cultures suggest that information is more optimally transmitted and stored as a result of neuronal networks being tuned near criticality [5, 28]. Therefore, criticality has been proposed as a set-point for the self-organization of cortical networks [29]. Nevertheless, some theoretical works have proposed that criticality only benefits the performance in complex cognitive tasks, while resting state conditions are less likely to benefit from these network dynamics [30]. Further support for this view identifies that healthy adults undertaking working memory and cognitive tasks have reported power-law scaling of response time fluctuation [31, 32]. Further, some forms of neurological dysfunction have been ascribed to impairment of critical dynamics [33–35]. While indicative, it has also been recognised that power laws are insufficient to infer criticality, since they can emerge from noise [36]. Further findings identifying linkages between criticality and stimulus discrimination, attention, language acquisition, fluid intelligence, and even conscious (awake) behaviour further complicate interpretations [37–43].

Consequently, there is still a lack of experimental evidence demonstrating whether criticality is a general property of biological neuronal networks, possibly generated by homeostatic mechanisms, or whether it is related to the brain’s response to mere informational load, or a more complex association with cognition. A question that still remains to be answered is whether cortical neuronal networks display a near-critical state during spontaneous activity or whether they only display near-critical states with structured information input - which for *in vivo* processing would typically occur when undertaking cognitive processing. An additional concern here is that functionally defined neural networks are rarely isolated from the many other connected networks of the intact brain, making it difficult to discern truly local critical functional dynamics as opposed to patterns derived from other regions [44].

To better evaluate these questions on the role of neural criticality, data was analysed from an *in vitro* neural network of cortical neurons which was trained to play the game ‘Pong’. We utilized *DishBrain*, a novel system shown to display goal-directed activity changes by harnessing the inherent adaptive computation of neurons [45]. We hypothesise that near-critical network behavior emerges when neural networks receive structured sensory input, and that this system would develop a network structure closer to critical states with successful task acquisition.

## Results

Cortical cells, either differentiated from human induced pluripotent stem cells (hiPSC) or derived from E15 mouse embryos, were subjected to the gameplay and rest conditions in the *DishBrain* system as previously described[45]. Hit-to-miss ratio and distance from the critical state were compared in different experimental conditions - see Supplementary Information. The measurements were carried out in both the conditions of (i) *Gameplay*, where cells adjusted paddle position through activity changes and received information about the position of the ball and the closed-loop response to their control of it, and (ii) *Rest*, where neuronal activity adjusted the paddle position, but received no input, in order to give a matched control. For more details, see Supplementary Information.

Neuronal avalanches were identified in network recordings. The scale-free dynamics of detected neuronal avalanches, as well as the Deviation from Criticality Coefficient (DCC), Branching Ratio (BR), and Shape Collapse error (SC error) were evaluated to identify whether the recordings were tuned near criticality. Figures 1.a-f provide a visual overview of the framework utilized in this study to investigate how far the dynamics of *in vitro* networks of cortical neurons are from criticality and whether this distance can accurately distinguish between task-present and task-absent states being processed by the neurons. Figure 1.i summarises these metrics of criticality and their formulations (see Supplementary Information: A.6 [3, 46, 47]). At criticality, BR of the network is tuned near 1.0 while DCC and SC error diminish to 0.

**Figure 1:**
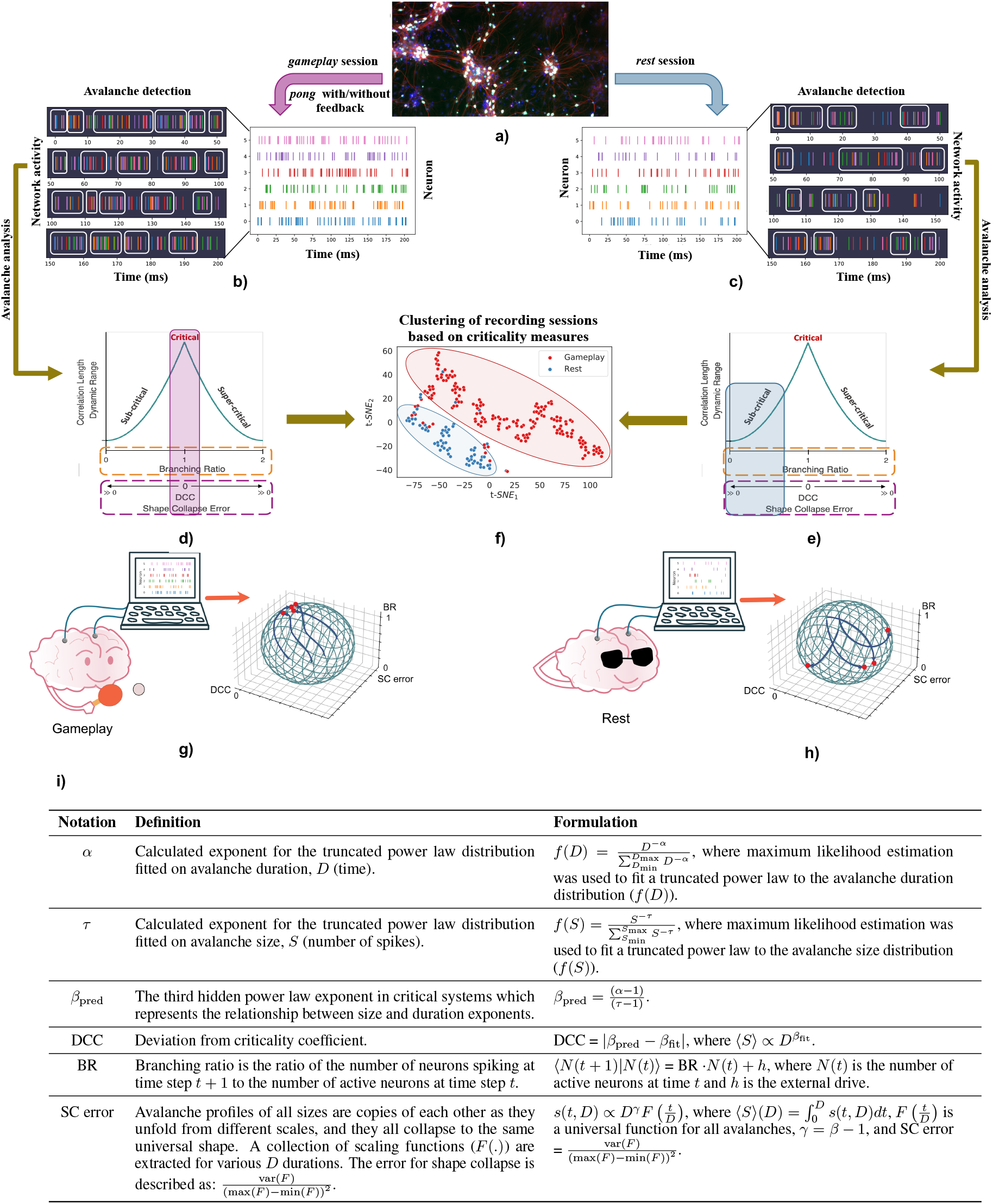
Schematic overview of study: **a)** Showing cortical cells harvested from embryonic rodents. **b, c)** The recorded population activity from these cortical cells is then binned to 50 ms bins during both *Gameplay* and *Rest* sessions. The neuronal avalanches are cascades of network activity that surpass a certain activity threshold for a certain duration of time, which are then extracted by bin. **d, e)** Avalanches are utilized to examine the criticality metrics in the neuronal network’s activity patterns to identify the working regime of each recording in terms of being sub-, super-, or near-critical. **f)** The same measures of criticality are used to cluster the recordings between two groups of *Gameplay* and *Rest*. **g, h)** Illustration of the experimental pipeline in which cultured cortical networks are recorded during *gameplay* and *rest* states. The recorded neuronal activities are then employed to extract the 3 metrics of criticality (namely Branching Ratio (BR), Deviation from Criticality Coefficient (DCC), and Shape Collapse error (SC error)) which are found to move towards the critical point during *Gameplay* (g) and move further from that point during *Rest* (h). **i)** Criticality Parameters and Metrics with details of their formulation.

### Cultured Cortical Networks Show Markers of Criticality When Engaged in a Task but Not When Resting

Data from 14 different cultures integrated on HD-MEAs during 308 experimental (192 *Gameplay*; 116 *Rest*) sessions were recorded and discretized into 50 ms bins. The sum of activities from all the recording channels in each time bin denotes the network activity. The network state was then evaluated using each of the described measures of criticality. Figure 2 illustrates the fitted PDF functions to avalanche size and duration and the associated pair of exponents (*τ* and *α*); the exponent is the slope of the line in a log-log plot. The associated DCCs extracted from the network are also represented. Two sample cultures were utilized to illustrate the comparison between the network’s dynamical state during *Rest* and *Gameplay*. Fitted power law distributions, DCC values, and the span of distributions in both size and duration domains are visualized in a *Rest* session (e.g., Session 1) against a *Gameplay* session (e.g. Session 4.)

**Figure 2:**
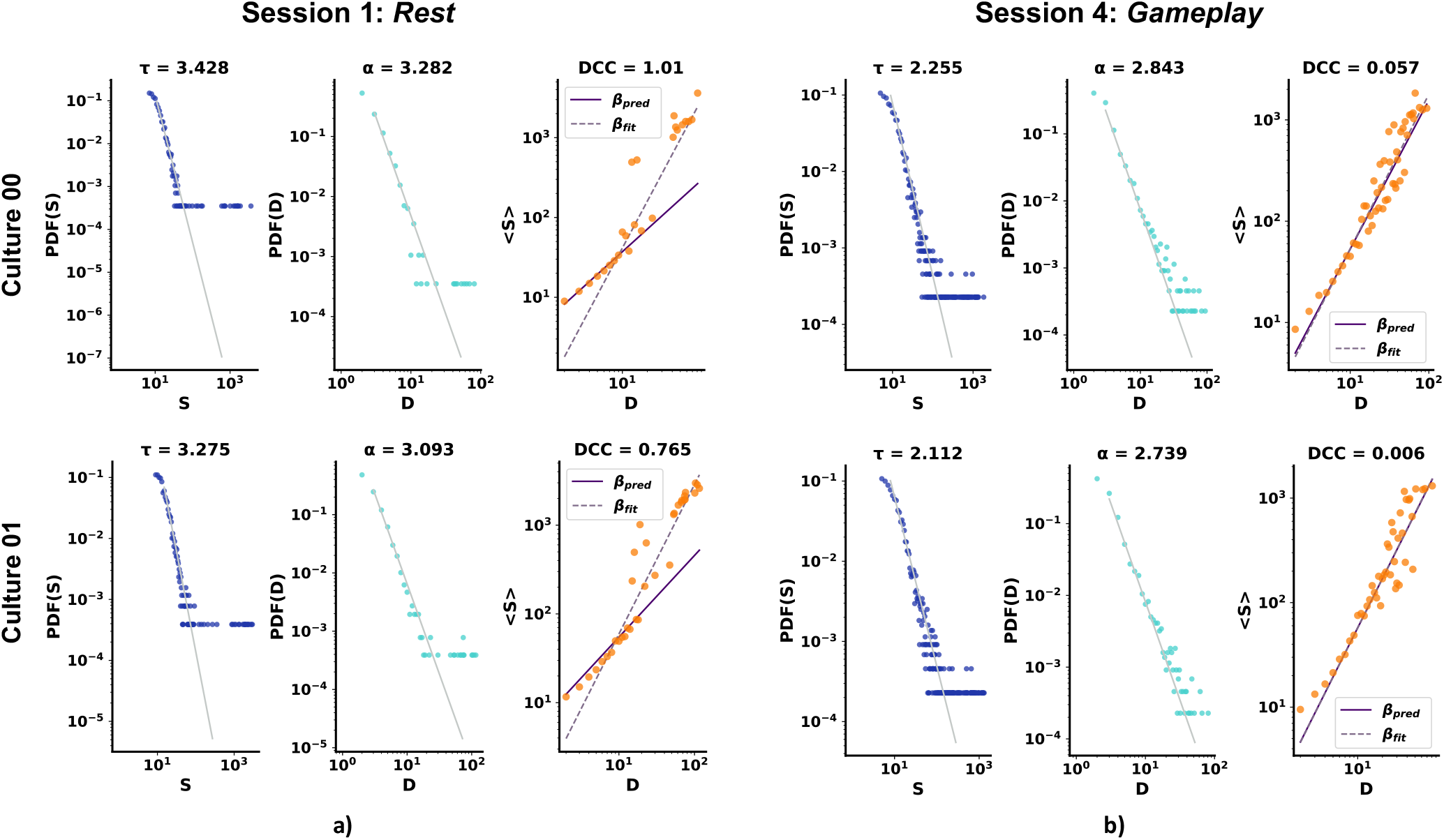
Avalanche size and duration PDF plots and the calculated DCC values for 2 sample cortical cultures at **a)** *Rest* (i.e. Session 1) and **b)** *Gameplay* (i.e. Session 4) of the same experiment and the corresponding *α* and *τ* exponents. DCC is given by |*β*_pred_ *-β*_fit_|; for details, see Table 1.a and A.8.

Additionally, BR and SC error were also extracted for all cultures in recording sessions 0 to 4.

Figure 3.a-c illustrate a general comparison between the critical and non-critical dynamics in terms of each of the introduced criticality metrics, DCC (3.a), BR (3.b), and SC error (3.c). An Alexander-Govern approximation test was run to investigate the significance of the differences between the two groups for each extracted metric. Figure 3.d-f illustrate the distribution of the criticality metrics in different recording sessions of the experiments. Comparison of the *Rest* (colored in teal) and *Gameplay* (colored in pink) sessions indicates the shift of cultured cortical network dynamics towards criticality during the task-present sessions. The *Gameplay* “Hit to Miss Ratio” (H/M ratio) - number of accurate “hits” to the number of “missed” balls - was also found to be significantly higher than during *Rest*. A summary of the statistical comparisons including the comparison of H/M ratio is given in Figure 3.g.

**Figure 3:**
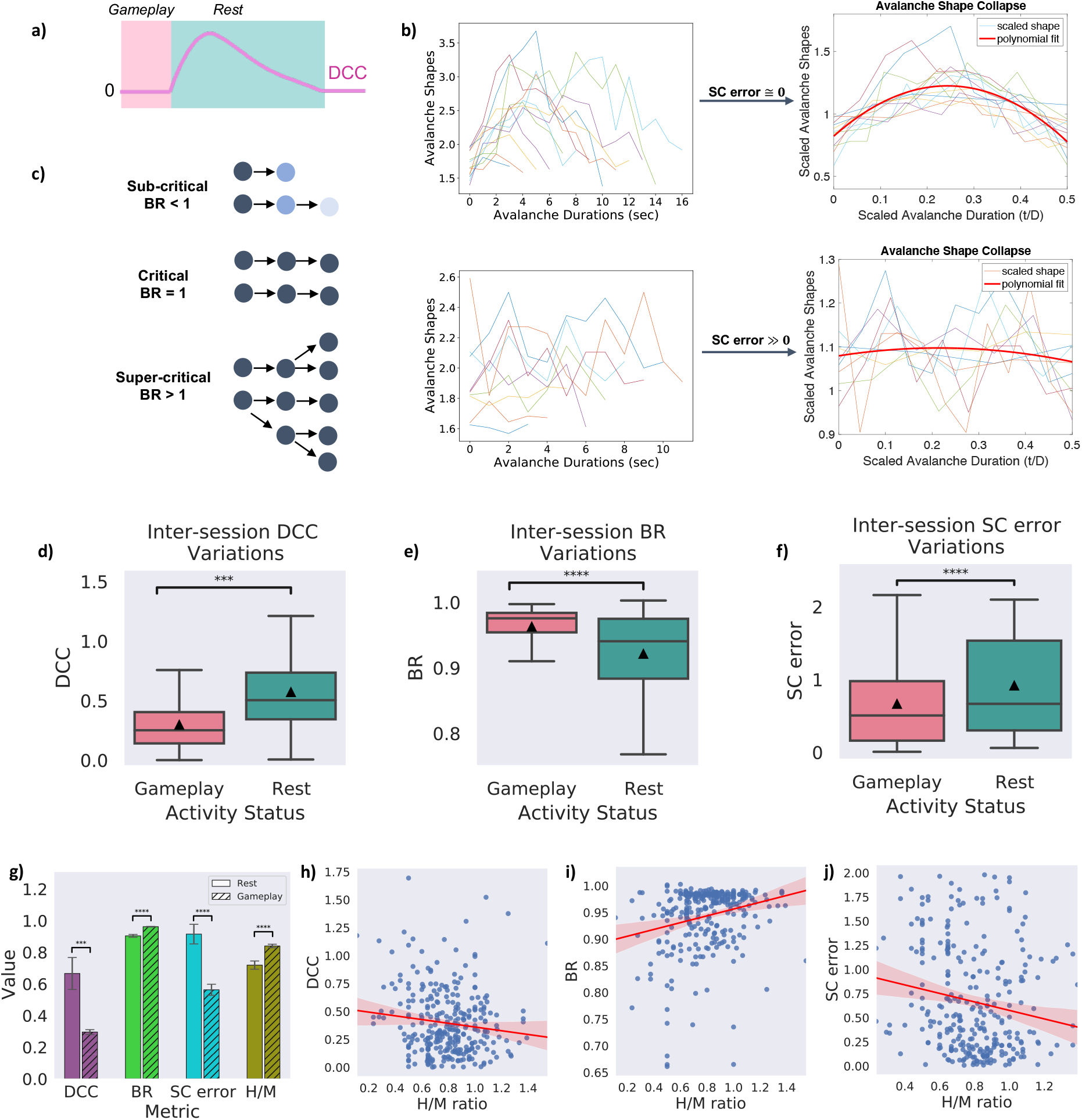
Comparison between critical and non-critical dynamics: **a)** Illustration of the course of the change expected to be observed in the DCC measure when transitioning between near-critical and non-critical regimes. **b)** Comparison of the shape collapse error while scaling avalanche shapes in 2 sample recording sessions. Scaled avalanches across a range of durations show little error around the polynomial fit in the upper row (indicative of a near-critical regime) while this error increases significantly in the data represented at the bottom row (indicative of a non-critical regime). **c)** Effect of branching ratio (BR) on activity propagation through a network over time. In critical regimes, BR = 1.0 and, on average, activity neither saturates nor decays across time. **d, e, f)** DCC, BR, and SC error extracted for all the recordings and compared between *Gameplay* and *Rest*. The illustrated trend in all measures supports the conclusion of the system tuning near criticality during *Gameplay*. The *Gameplay* recordings display DCC and SC error values closer to 0 and branching ratios closer to 1; features which are missing in the *Rest* recordings. Box plots show interquartile range, with bars demonstrating 1.5X interquartile range, the line marks the median and ▴ marks the mean. Error bands, 1 SE. **g)** Summary of the key characteristics of a critical system compared between all *Rest* and *Gameplay* sessions as well as the corresponding performance level in terms of the observed H/M ratio. ***p < 5 × 10^−4^, ****p < 5 × 10^−5^. Error bars, SEM. **h)** A weakly significant negative correlation was found between DCC and the neuronal culture performance in terms of H/M ratio (r = −0.13, p < 0.05, Pearson Correlation test). **i)** A strongly significant positive association was observed between BR and H/M ratio (r = 0.24, p < 0.00005, Pearson Correlation test), and **j)** A strongly significant negative correlations was found between SC error and H/M ratio (r = −0.17, p < 0.005, Pearson Correlation test).

These results indicate the shift towards self-organized criticality of the neural cultures in these experiments when exposed to external structured information such as the game environment of ‘Pong’. In contrast, cultured cortical networks deviated from the critical state during rest sessions when the paddle was solely affected by the neurons’ spontaneous activities. When the cells were not presented any external information about the status of the ball or the game, the network parameters indicated a sub-critical system. These results suggest that during task-present conditions (here accompanied by learning, which is reflected in the improved H/M ratio of experimental cultures), the cultured cortical network tunes itself near criticality.

Notably, deviation from criticality was also measured in time shuffled data acquired from the *Gameplay* sessions. These data preserved the spatial correlations but randomized the temporal structure and obtained a significantly higher DCC value compared to the original data, indicating a larger deviation from criticality compared to the original data (DCC for time shuffled and original recordings: 0.627 *±* 0.087 and 0.296 *±* 0.015 respectively, p< 0.0005, Alexander-Govern approximation test). Specifically, we sought further control data by analysing time shuffled data to detect the sensitivity to the ensemble activity’s temporal structure. This control is important since the comparison can then eliminate the potential role of temporal random effects in detecting the critical dynamics.

Furthermore, to determine whether the identified criticality metrics correlated with game performance, exploratory uncorrected Pearson’s correlations were computed for criticality metrics and H/M ratio for all recording sessions (see Figure 3.h-j). While significant negative correlation was found between DCC and H/M ratio (r = −0.13, p < 0.05, Pearson Correlation test) as well as SC error and H/M ratio (r = −0.17, p < 0.005, Pearson Correlation test), a strong positive association was observed between BR and culture performance represented by H/M ratio (r = 0.24, p < 0.00005, Pearson Correlation test). This indicates that network dynamics closer to criticality may be related to better performance.

### Culture *Gameplay* vs *Rest* Status is Predicted by Criticality Metrics and H/M ratio

Binary classification of the data was performed to predict group membership of each recording session and assign it to either the *Rest* or *Gameplay* classes. Three different classification algorithms were utilized: Logistic Regression, Support Vector Machines (SVM), and Random Forests. Figure 4.a represents the mean prediction accuracy for various classification methods as well as different approaches in assigning feature vectors to the data points. 4-Metrics refers to the case where a 4 dimensional vector of all the 4 metrics represented in Figure 3.g were used to represent each data point. 3-Criticality metrics indicates a case where only 3 criticality metrics are used to form the feature vectors. The conditions where each metric is separately used to represent the data points is also included. The results demonstrate that the highest accuracy of prediction can be achieved using all 4 criticality metrics accompanied by the culture’s H/M ratio. Nevertheless, it was also found that merely employing the criticality measures is sufficient for an accurate prediction (up to 92.41%) of the culture’s status in terms of it being task-present or task-absent (i.e., the default resting state). These findings suggest that knowledge about a neuronal network’s distance from criticality may be adequate for distinguishing between task-present and task-absent states and whether the input information is being optimally processed. Data representations were visualized using the obtained feature vectors. Since the 4-Metrics representation proved to be the most effective representation given the results in the table of Figure 4.a, we considered this case for the visualization task. A standard t-SNE algorithm [48] visualized the data representations as per Figure 4.b. A 2-dimensional visualization of the sessions is obtained with each colored dot as an indicator of each data point. The Kullback-Leibler divergence indicating the error between the pairwise similarities of the input and their corresponding projections in the resulting 2-dimensional mapping is 0.373, indicating an accurate network representation.

**Figure 4:**
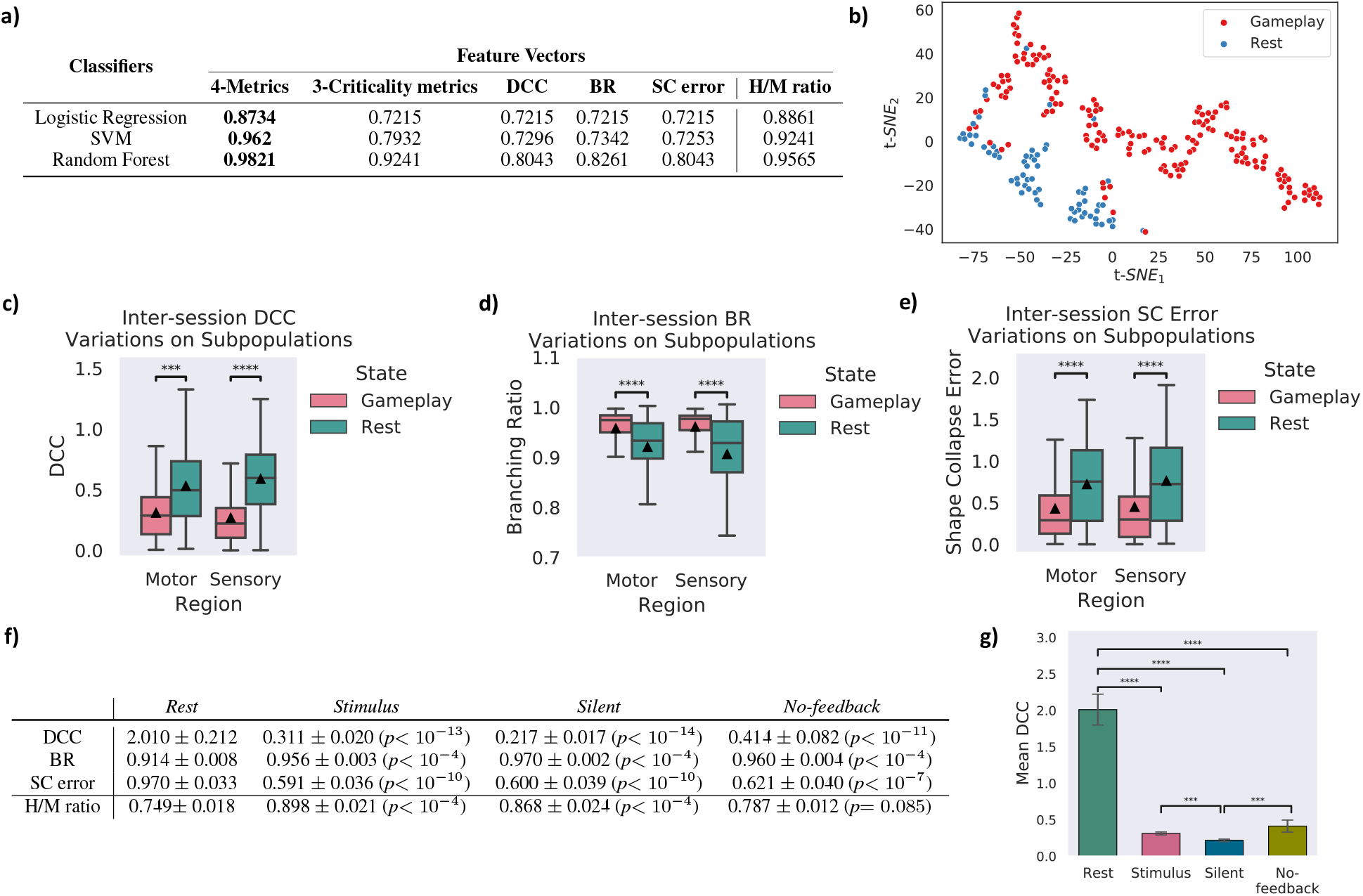
**a)** Comparison of the mean prediction accuracy for different binary classifiers on all the recorded sessions. **b)** Visualization of the extracted representation for each data point using the t-SNE algorithm in a 2-dimensional space (i.e. dimensions t-SNE_1_ and t-SNE_2_). The two *Rest* and *Gameplay* classes are illustrated with different colors. **c)** DCC, **d)** BR, and **e)** SC error variations between *Rest* and *Gameplay* sessions in separate motor and sensory regions of the cultures. The illustrated trend in all three measures on the subpopulations is in line with the previous conclusion about the entire population. The similar pattern in these results also states that during *Gameplay* the neuronal ensembles move near criticality while in *Rest*, they are further from a critical state. ***p < 10^−3^, ****p < 10^−5^. Box plots show interquartile range, with bars demonstrating 1.5X interquartile range, the line marks the median and ▴ marks the mean. Error bands, 1 SE. **f)** Comparison of the 4 extracted criticality measure as well as the H/M ratio for the *Rest* and *Gameplay* groups. The *p-*value of an Alexander-Govern approximation test is reported for each measure in comparison of each feedback condition with *Rest* sessions. **g)** Comparing the average DCC measure calculated in different feedback conditions with the *Rest* sessions. ***p < 5 × 10^3^, ****p < 10^−10^. Error bars, SEM.

### Motor and Sensory Subpopulations Inherit the Criticality Characteristics of the Entire Neuronal Ensemble

In the *DishBrain* system’ configuration, a specific frequency and voltage are applied to key electrodes in the predefined sensory areas, as described in [45]. Then, different predefined motor region configurations are examined to select the configuration that maximises performance. The paddle moves in a corresponding direction based on the region with the higher activity.

We evaluated these predefined regions and the overlaid neuronal subpopulations in terms of their activity dynamics. Consequently, the introduced criticality metrics were measured from the recorded activities in each of these subpopulations separately. Figure 4.c-e demonstrates that these subpopulations also exhibit similar features of a near-critical system when exposed to the *Gameplay* setting. Yet both motor and sensory neuronal populations were identified to be statistically significantly closer to criticality compared to *Rest* session recordings from the same neuronal subpopulations.

### Feedback Is Required for Improved Game Performance in a Critical System

Biological neuronal networks typically require feedback for learning to occur - i.e., a closed-loop between action and consequence. In a closed-loop system, feedback is provided on the causal effects of the neuronal culture’s behavior Three different feedback conditions were employed in this study. Condition 1 (*Stimulus*), is where predictable and unpredictable stimulus are administered when the cultures behaved desirably or not, respectively (results reported previously). Condition 2 (*Silent*), involves the above stimulus feedback being replaced with a matching time-period where all stimulation was withheld. Condition 3 (*No-feedback*), involves a more drastic change to the gameplay environment, where the ability for the ball to be missed was removed, so when the paddle failed to intercept the ball, the ball bounced instead of triggering a reset, and the game continued uninterrupted [45] (see Supplementary Information: A.4). Performance of the cultures in terms of their H/M ratio, as well as their criticality characteristics were measured under all three feedback conditions and then compared to the *Rest* sessions. Table 4.f represent the results (Mean *±* SE) for 14, 15, and 12 different cultures under *Stimulus, Silent*, and *No-feedback* conditions respectively. Overall 113, 117, and 95 sessions were recorded under the different feedback types and were compared to 209 *Rest* sessions obtained from the total of 41 cultures under experiment. The reported p−values represent the significance of the difference between the obtained measures in each feedback condition and the *Rest* cultures. It is very interesting to observe that the *Silent* condition shows significant performance in the game (H/M ratio) as well as showing dynamical features that are indicative of a near-critical system. The deviation from criticality (DCC) is significantly lower in the *Silent* condition compared to *Stimulus* or *No-feedback* (p < 0.005, Alexander-Govern approximation test) conditions (see Figure 4.g. This difference was not significant when comparing *Stimulus* and *No-feedback* conditions. While comparing the gameplay characteristics of the cultures (H/M ratio), the *Stimulus* and *Silent* conditions both significantly outperform the *No-feedback* conditions (p <0.0005 and p <0.005, Alexander-Govern approximation test). While the *No-feedback* system also represents features characterizing a near-critical dynamics (although to a lesser extent compared to the other two closed-loop systems), the game performance is significantly deteriorated in this case (no significant outperformance compared to the *Rest* state, p = 0.085, Alexander-Govern approximation test). This demonstrates that fine tuning near criticality may be necessary for optimal information processing when facing an increased load. Nonetheless, criticality may not be sufficient for a neuronal network to achieve its learning and memory goals in the absence of additional information regarding the consequences of previous actions, i.e., feedback.

## Discussion

Near-critical dynamics in the brain remain a fascinating phenomenon and here we demonstrate it is readily observable for *in vitro* neuronal cultures when embodied in a virtual environment [45] through the structured stimulation. While novel, this finding accords with increasing evidence linking near-critical dynamics in the brain with cognitive behaviour [49–51].Yet, by utilizing *in vitro* cortical networks integrated with *in silico* computing via HD-MEA to experimentally explore the notion of criticality under task-present and task-absent states some key tentative conclusions can be made. As evidenced through multiple features expected of a near critical system, we found that cultured networks of cortical neurons self-organized to display these key markers when undertaking task acquisition but not when unstimulated. Through this it was robustly observed that *in vitro* cortical neurons exhibited markers of criticality when actively engaged in a task and receiving feedback contingent on neuronal activity modulating the stimulated world. While many previous studies rely on identifying power-law scaling in temporal and partial domains [31, 32], power laws may emerge from noise [36]. Generally, power laws should be accompanied by independent stochastic surrogates, such as disconnected nodes in a complex system [36].As such, we co-analysed the described three established markers of criticality on spiking data generated in this system [5, 46]. We observed an exceptionally high degree of qualitative concordance between these different measures, adding confidence to the internal validity of the results. The current study adds weight to previous findings in literature. It has been proposed that certain features of learning, including optimal information capacity and transmission are optimized at criticality [52]. Indeed, many studies have recently identified *in vivo* that cortical networks typically function near a critical point [4, 5], the extent of which shows key correlations with performance [3]. Therefore, one interpretation of our finding that *in vitro* neuronal networks show markers of criticality only when presented structured information, is that this activity arises during information processing. In contrast, while in the default resting state of *in vitro* neuronal networks, i.e., not embodied within a game environment, despite spontaneous activity exhibiting neuronal avalanches, cultures no longer display hallmarks of criticality. The extent of this difference based on markers of criticality alone was stark enough to predict whether a given culture was actively engaged in gameplay or resting with a 92.41% accuracy. When performance data was included this accuracy increased to 98.21%, further supporting the dramatic difference between resting and active cultures. The additional finding that these markers of criticality were persistent across sub-populations defined by their external relationship to the game-world for the neuronal cultures, suggests a network wide coordination of activity. Taken in concert with past research identifying power-law like behaviour in humans undergoing cognitive tasks, this is indicative of a network wide fundamental computation underlying information processing which may be ongoing in these cultures only when actively engaged in a task or otherwise presented structured information [31, 32].

However, these results, although in agreement with some studies, must be taken in a wider context. For example, the importance of critical state dynamics in language acquisition has been highlighted [39, 40]. Moreover, a recent resting-state fMRI study of neurotypical adults with varying IQs have found a connection between high fluid intelligence and close proximity to a critical state in a spin-glass model [37]. In addition, the conscious states of mind have been linked to near-critical slow cortical electrodynamics, suggesting that the disruptions in information processing during unconscious states is due to the transition of low-frequency cortical electric oscillations away from the critical point [38]. In contrast, several studies demonstrate more ambiguous results around the relationship between electrical brain response to increased cognitive load [53–56]. Furthermore, in the majority of previous studies, the observation of power laws was regarded as the primary indicator of criticality. As noted above, the criticisms of this approach [46, 57, 58] render this data insufficient to robustly establish that a system is certainly operating at criticality.

The ambiguity around the role of criticality in human cognition prevents any robust conclusion that the evidence of criticality in these cultures while actively engaged in gameplay is prima facia evidence of network wide learning and/or cognition. A more nuanced perspective is that criticality is not tied to general processing, learning or cognition, but is rather optimised for specific tasks or types of information processing. Previously, criticality was found to be linked to stimulus discrimination, yet decreased stimulus detection [41]. This nuanced view is supported by the observed variation in mean DCC between different feedback types for when the neuronal networks were engaged in *gameplay*. Most notably, the finding that the open-loop no-feedback condition, where modulated activity from the culture was unable to affect the game outcome or alter the feedback received, showed considerably closer dynamics to criticality than when cultures were resting is interesting. This suggests that structured information input alone may be sufficient to induce these near-critical states in neuronal systems, however, information alone is not sufficient in creating an evolving learning system as feedback is required as well. Furthermore, feedback does not necessarily need to be a positive addition to the system as identified in the experiments utilizing *Silent* feedback conditions. Taken in the context of attentional engagement and criticality, it is possible that an external source of information is necessary to drive these characteristics observed here [42, 43]. // Nonetheless, allowing cultures to be embodied and alter the environmental stimulation through action does seem to push cultures closer to a critical dynamic, highlighting the need to consider the notion of criticality as a spectrum. What this research does robustly demonstrate is one end of this spectrum: criticality requires the input of structured information to a system to arise. This finding is entirely consistent with all rigorous studies into criticality. Future work is still needed to further explain the more specific role of criticality in information processing and cognition - both *in vivo* and *in vitro*. Yet while questions about how neural criticality is linked with human cognition remain, ultimately, this work has suitably established that closeness to criticality appears as a fundamental property to neuronal assemblies and is influenced by the input of structured information. This provides a compelling pathway to better investigate the critical aspects of how our brains process information.

## Acknowledgements

F.H. was supported by the Melbourne Research Scholarship. A.N.B. was supported by the Australian Government through the Australian Research Council’s Discovery Projects funding scheme (Project DP220101166). C.F. was supported by the RMH Neuroscience Foundation.

## Author Contributions

Conceptualization, B.J.K., F.H., A.B., C.F.; methodology, C.F., B.J.K., F.H., A.B.; software, F.H., B.J.K.; analysis, F.H., B.J.K.; cell culture, B.J.K.; data curation, B.J.K., F.H.; writing—original draft preparation, F.H., B.J.K.; writing—review and editing, F.H., B.J.K., C.F., A.B., D.D.; visualization, F.H., B.J.K., D.D.; project administration, C.F., B.J.K., A.B.; supervision, C.F., B.J.K., A.B.

## Declaration Of Interests

B.J.K. & D.D. are employees of Cortical Labs. B.J.K. is a shareholder of Cortical Labs and holds an interest in patents related to this publication. F.H. received funding from Cortical Labs for work related to this publication.

## Notes

### Competing Interest Statement

The authors have declared no competing interest.

