## Supplemental Materials for "Critical dynamics arise during structured information presentation: analysis of embodied *in vitro* neuronal networks"

### A Supplementary Information

#### A.1 Cell Culture

Neural cells were cultured either from the cortices of E15.5 mouse embryos or differentiated from human induced pluripotent stem cells via a dual SMAD inhibition (DSI) protocol or through a lentivirus based NGN2 direct differentation protocols as previously described [45]. Cells were cultured until plating. For primary mouse neurons this occurred at day-in-vitro (DIV) 0, for DSI cultures this occurred at between DIV 30 - 33 depending culture development, for NGN2 cultures this occured at DIV 3.

#### A.2 MEA Setup and Plating

MaxOne Multielectrode Arrays (MEA; Maxwell Biosystems, AG, Switzerland) was used and is a high-resolution electrophysiology platform featuring 26,000 platinum electrodes arranged over an $8mm^2$ surface. The MaxOne system is based on complementary meta-oxide-semiconductor (CMOS) technology and allows recording from up to 1024 channels. MEAs were coated with either polyethylenimine (PEI) in borate buffer for primary culture cells or Poly-D-Lysine for cells from an iPSC background before being coated with either 10 µg/ml mouse laminin or 10 µg/ml human 521 Laminin (Stemcell Technologies Australia, Melbourne, Australia) respectively to facilitate cell adhesion. Approximately $10^6$ cells were plated on MEA after preparation as per [45]. Cells were allowed approximately one hour to adhere to MEA surface before the well was flooded. The day after plating, cell culture media was changed for all culture types to BrainPhys™ Neuronal Medium (Stemcell Technologies Australia, Melbourne, Australia) supplemented with 1% penicillin-streptomycin. Cultures were maintained in a low O2 incubator kept at 5% CO2, 5% O2, 36°C and 80% relative humidity. Every two days, half the media from each well was removed and replaced with free media. Media changes always occurred after all recording sessions.

#### A.3 Dishbrain platform and electrode configuration

The current DishBrain platform is configured as a low-latency, real-time MEA control system with on-line spike detection and recording software. The DishBrain platform provides on-line spike detection and recording configured as a low-latency, real-time MEA control. The DishBrain software runs at 20 kHz and allows recording at an incredibly fine timescale. There is the option of recording spikes in binary files, and regardless of recording, they are counted over a period of 10 milliseconds (200 samples), at which point the game environment is provided with how many spikes are detected in each electrode in each predefined motor region as described below. Based on which motor region the spikes occurred in, they are interpreted as motor activity, moving the 'paddle' up or down in the virtual space. As the ball moves around the play area at a fixed speed and bounces off the edge of the play area and the paddle, the pong game is also updated at every 10ms interval. Once the ball hits the edge of the play area behind the paddle, one rally of pong has come to an end. The game environment will instead determine which type of feedback to apply at the end of the rally: random, silent, or none. Feedback is also provided when the ball contacts the paddle under the standard stimulus condition. A 'stimulation sequencer' module tracks the location of the ball relative to the paddle during each rally and encodes it as stimulation to one of eight stimulation sites. Each time a sample is received from the MEA, the stimulation sequencer is updated 20,000 times a second, and after the previous lot of MEA commands has completed, it constructs a new sequence of MEA commands based on the information it has been configured to transmit based on both place codes and rate codes. The stimulations take the form of a short square bi-phasic pulse that is a positive voltage, then a negative voltage. This pulse sequence is read and applied to the electrode by a Digital to Analog Converter (or DAC) on the MEA. A real-time interactive version of the game visualiseris available at https://spikestream.corticallabs.com/. Alternatively, cells could be recorded at 'rest' in a gameplay environment where activity was recorded to move the paddle but no stimulation was delivered, with corresponding outcomes still recorded. Using this spontaneous activity alone as a baseline, the gameplay characteristics of a culture were determined. Low level code for interacting with Maxwell API was written in C to minimize processing latencies-so packet processing latency was typically $<50$ $\mu$s. High-level code was written in Python, including configuration setups and general instructions for game settings. A 5 ms spike-to-stim latency was achieved, which was substantially due to MaxOne's inflexible hardware buffering. Figure A1 illustrates a schematic view of Software components and data flow in the DishBrain closed loop system.

##### A.4 Input Configuration

As introduced in [45], stimulation is delivered at specific locations, frequency, and voltage to key electrodes in a topographically consistent manner in the sensory area relative to the current position of the paddle (Figure A4: Configuration 0). This mimics retinotopic and topographic representations found in many neural systems, which represent the external world [59, 60]. It was possible to deliver five types of input. Either the 'Sensory Stimulus' encoding the position of the ball, or one of four feedback protocols explained below: Unpredictable, Predictable, Silent, or No-feedback.

**Sensory Stimulus**: Due to the fact that neurons appeared robust to voltage stimulation, the voltage level was determined based on the evidence of neurological function. As a result, 75 mV was chosen as the sensory stimulation voltage to prevent forcing hyperpolarised cells to fire. This stimulation was applied to key electrodes relating to where the ball was relative to the paddle. Combining place coding with a rate coding, when the ball was closest to the opposing wall, stimuli were delivered at 4 Hz, increasing in a linear manner to 40 Hz when it reached the paddle wall.

**Unpredictable Stimulus**: For the standard stimulus feedback condition, cultures received unpredictable stimulation when they missed connecting the paddle with the 'ball', i.e. when a 'miss' occurred. Using a feedback stimulus at a voltage of 150 mV and a frequency of 5 Hz, unpredictable external stimulus could be added to the system. Random stimulation took place at random sites over the 8 predefined input electrodes at random timescales for a period of four seconds, followed by a configurable rest period of four seconds where stimulation paused, then the next rally began. In theory, the higher voltage than used for the Sensory Stimulus would force action potentials regardless of the state in which the cell was in, causing even greater disruption.

**Predictable Stimulus**: For the standard stimulus feedback condition, cultures were exposed to predictable stimulation

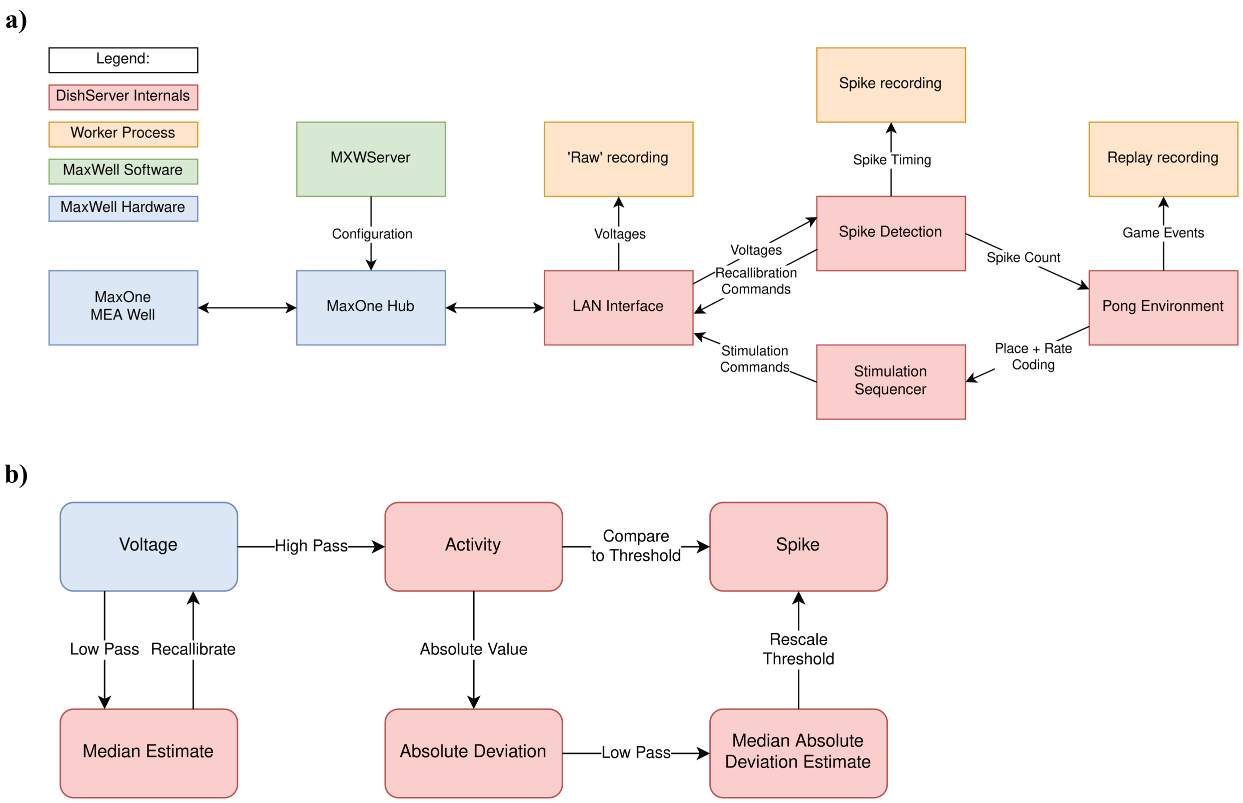


**Figure A1: a, b)** Schematics of software used for DishBrain. **a)** Software components and data flow in the DishBrain closed loop system. Voltage samples flow from the MEA to the 'Pong' environment, and sensory information flows from the 'Pong' environment back to the MEA, forming a closed loop. The blue rectangles mark proprietary pieces of hardware from MaxWell, including the MEA well which may contain a live culture of neurons. The green MXWServer is a piece of software provided by MaxWell which is used to configure the MEA and Hub, using a private API directly over the network. The red rectangles mark components of the 'DishServer' program, a high-performance program consisting of four components designed to run asynchronously, despite being run on a single CPU thread. The 'LAN Interface' component stores network state, for talking to the Hub, and produces arrays of voltage values for processing. Voltage values are passed to the 'Spike Detection' component, which stores feedback values and spike counts, and passes recalibration commands back to the LAN Interface. When the pong environment is ready to run, it updates the state of the paddle based on the spike counts, updates the state of the ball based on its velocity and collision conditions, and reconfigures the stimulation sequencer based on the relative position of the ball and current state of the game. The stimulation sequencer stores and updates indices and countdowns relating to the stimulations it must produce and converts these into commands each time the corresponding countdown reaches zero, which are finally passed back to the LAN Interface, to send to the MEA system, closing the loop. The procedures associated with each component are run one after the other in a simple loop control flow, but the 'Pong' environment only moves forward every 200th update, short-circuiting otherwise. Additionally, up to three worker processes are launched in parallel, depending on which parts of the system need to be recorded. They receive data from the main thread via shared memory and write it to file, allowing the main thread to continue processing data without having to hand control to the operating system and back again. **b)** Numeric operations in the real-time spike detection component of the DishBrain closed loop system, including multiple IIR filters. Running a virtual environment in a closed loop imposes strict performance requirements, and digital signal processing is the main bottleneck of this system, with close to 42 MB of data to process every second. Simple sequences of IIR digital filters is applied to incoming data, storing multiple arrays of 1024 feedback values in between each sample. First, spikes on the incoming data are detected by applying a high pass filter to determine the deviation of the activity, and comparing that to the MAD, which is itself calculated with a subsequent low pass filter. Then, a low pass filter is applied to the original data to determine whether the MEA hardware needs to be re-calibrated, affecting future samples. This system was able to keep up with the incoming data on a single thread of an Intel Core i7-8809G. Figures adapted from [45].

when a 'hit' was registered - that is, when the 'paddle' connected successfully with the 'ball'. This was delivered at 75mV at 100Hz over 100ms. This occurred when the simulated ball struck the paddle and replaced other sensory information for 100 ms. All 8 stimulation electrodes simultaneously received predictable stimulation at this frequency and period.

**Silent Feedback**: During Silent feedback period, the Unpredictable Stimulus described above was replaced with no stimulation for the same length of time. Predictable Stimulus feedback was also removed during Silent Feedback sessions. There is still a difference between this feedback and No-Feedback described below. This feedback is associated with culture activity in a closed-loop manner and therefore constitutes feedback.

**No-feedback:** As an open-loop condition, this assessment was designed to determine whether sensory stimulation is sufficient to drive learning in cultures. In other words, there was no feedback of any kind provided to the cultures based on their actions or outcomes. As described above, the cultures were given the same sensory stimulation as described above, and the outcome was determined by the same metric. However, when a ‘miss’ would otherwise occur, here instead the ball bounced off the wall behind the paddle, continuing the same trajectory – still recorded as a ‘miss’. This would otherwise end the rally. The ‘ball’ would be recorded as a ‘hit’ whenever it connected with the simulated paddle. So, under No-Feedback, the entire gameplay session is essentially one rally in which the simulated ball’s final position can be predicted from its initial vector, but scoring occurs normally as usual.

Figure A2 represents a schematic showing the different phases of stimulation and the information delivery to the culture while Figure A3 demonstrates different simulated gameplay environments employing different feedback protocols as above.

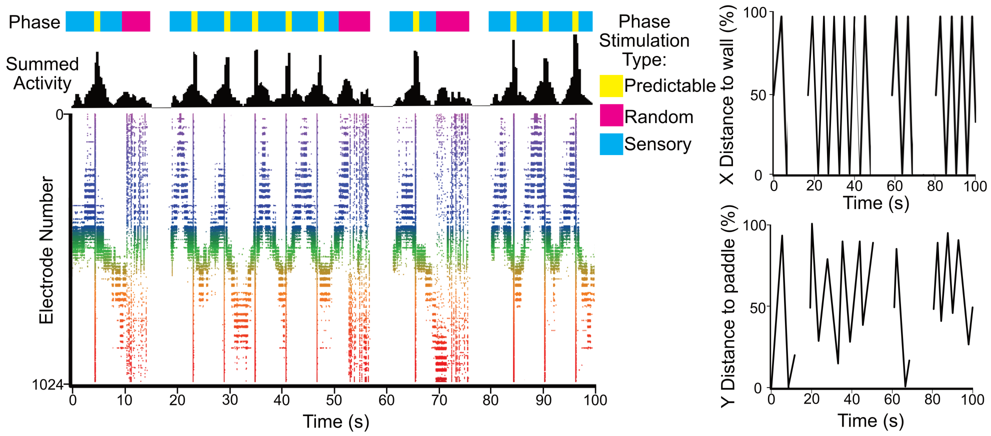


**Figure A2:** Presents a schematic showing the different phases of stimulation which provides information about the environment to the culture, in line with this is the corresponding input voltage and how that voltage appears on the raster plot over 100 seconds. The appearance of random stimulation after a ball missing vs system wide predictable stimulation upon a successful hit is apparent across all three representations. This corresponds to the images on the right which show the position of the ball on both x and y axis relative to the paddle and backwall in % of total distance is shown on the same timescale. Figure adapted from [45].

##### A.5 Output Configuration

A total of 1024 electrodes were routed on the HD-MEA to record activity. The ‘Sensory’ area, where stimulation electrodes were embedded as described above consisted of 626 electrodes. The remaining output electrodes were divided into predefined motor regionson the MEA, consisting of four regions that were defined either as motor region 1 or motor region 2 as shown in Figure A4. Since it was technically difficult to cultivate neurons that displayed perfectly symmetrical activity in both these regions, ’gain’ was added to the system. These took a real-time value based on

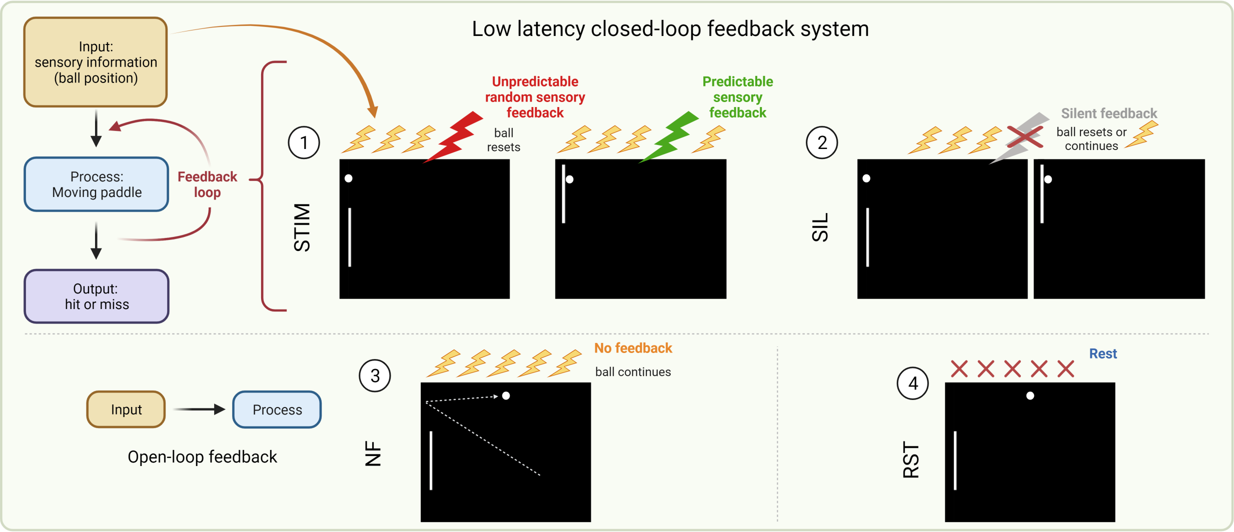


**Figure A3:** Different DishBrain environments were utilised to demonstrate: (1&2) low latency closed-loop feedback system (stimulation (STIM) & silent (SIL) treatment); (3) No feedback (NF) system to demonstrate an open-loop feedback configuration; and (4) rest (RST) configuration to demonstrate a system in which sensory information (yellow bolt) is absent. Figure adapted from [45]

.

the mean firing in each motor region and multiplied it to achieve a target value of 20 $Hz$ across the entire region. Consequently, changes in activity in each of the two regions could affect the position of the paddle, even if they displayed different latent spontaneous activity.

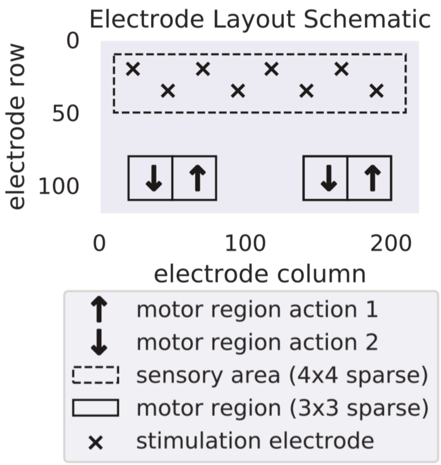


**Figure A4:** Electrode layout schematic for DishBrain Pong-world gameplay. Figure adapted from [45].

#### A.6 Data analysis

The avalanche analysis is performed in order to study the network in terms of its distance from criticality. The start and stop of an avalanche are determined by crossing a threshold of network activity [3]. An avalanche can be initiated by spikes from any and all neurons within a region of interest. The number of contributing spikes in each avalanche ($S$)

and the total duration of the event ($D$) are then measured. To demonstrate the distance from criticality in a cultured cortical network, the presence of the following markers were investigated in the dynamics of our data: 1) Power law observables; 2) Exponent relation; 3) Branching ratio parameter; and 4) Scaling function. Certain criteria on these markers are the necessary conditions of a critical regime and meeting those criteria can indicate with a high confidence whether the system lies near a critical point [5, 46].

##### A.7 Power law observables

A critical system has interacting components (here, neurons) that show some fluctuation in their activity while also maintaining a level of correlation between their individual activities (here, individual spiking). Criticality implies that the system is defined by scale free dynamics and that events in both the spatial and temporal domains obey power laws [57, 61, 62]. For the networks considered here, events are contiguous cascades of spiking activity, rather than limited local bursts of spiking activity or huge network-wide spiking events. These contiguous cascades of spiking activity are called *neuronal avalanches*.

To investigate this property in our BNN system, binary spike trains of each neuron's activity were utilized. The whole duration of each recording session was discretized to 50 ms bins. The sum of all cells' activities in each time bin was used as the network activity. Next, a threshold of 40% of the median spiking activity in the network among all time bins was introduced. The start and end points of an avalanche were defined as the time points when the network activity crossed this threshold value from below and then above [63]. Our results were statistically robust across a range of activity thresholds between 30% and 70%. The size of an avalanche, $S$, is the total number of spikes during the avalanche. The avalanche duration, $D$, is the time between threshold crossings. Similar to [3], maximum likelihood estimation was used to fit a truncated power law to the avalanche size distribution:

$$f(S) = \frac{S^{-\tau}}{\sum_{S_{\min}}^{S_{\max}} S^{-\tau}}, \tag{A.1}$$

where $\tau$ is the power law exponent corresponding to avalanche sizes. For a neuronal recording session in which $N_A$ avalanches are detected, the fitting process to obtain the above equation is the following iterative procedure [64]:

1. Find the maximum observed avalanche size $S_{\max}$.
2. Evaluate the three different power law exponents, $\tau$, for the 3 smallest avalanche sizes observed, $S_{\min}$.
3. Calculate the *Kolmogorov-Smirnov* (KS) test for this estimation to determine the goodness-of-fit between the fitted power law and the empirical distribution.
4. Among the obtained KS values, choose the smallest one, together with the corresponding $\tau$ and $S_{\min}$ values.
5. Complete the estimation if KS $< \frac{1}{\sqrt{N_A}}$ or otherwise repeat steps 2 to 5 with $S_{\max}$ reduced by 1 until this condition is met.

Steps 3 to 5 are necessary to ensure the data distribution indeed comes from a power law rather than another candidate heavy tailed distribution, such as log normal and stretched exponential forms [65]. Applying the exact same procedure to the set of $D$ avalanche events, the corresponding power law exponent of $\alpha$ was calculated for the entire avalanche duration distribution.

To test the validity of a power law fit to avalanche distributions, hypothesis testing was performed as described in [3]. For this purpose, the power law exponent, the number of detected avalanches, and the minimum and maximum avalanche sizes were set the same as the experimental avalanche distribution to generate 1000 artificial power law distributions. We generated these surrogate distributions using the inverse method as $S = S_{min}(1-r)^{\frac{-1}{\tau-1}}$ where $r$ was a random number sampled from a uniform distribution between 0 and 1. Then any surrogate distribution was upper-truncated at the maximum cut-off equivalent to $S_{max}$ from the empirical data. The KS statistics was then employed to estimate the distance between the simulated surrogate distributions and a perfect power law. The $p$ value determining the significance level was then equal to the ratio of the surrogate distributions with KS values smaller than the KS value of the corresponding experimental avalanche distribution. With significance level set to 0.05, $p < 0.05$ implies a rejection of the power law hypothesis while $p \geq 0.05$ suggests the power law hypothesis was not rejected (the fit was good).

##### A.8 Exponent relation and Deviation from Criticality Coefficient (DCC)

In critical systems, there is another exponent relationship between the power law parameters ($\alpha$ and $\tau$) and the exponent of mean avalanche sizes ($\langle S \rangle$), given their duration, $D$ [66]. We first find this third power law exponent of the system, $\beta$, from the experimental data using linear regression given the following exponent relation is present in a critical system:

$$\langle S \rangle \propto D^{\beta_{\text{fit}}}. \tag{A.2}$$

This third power law exponent also relates the size and duration distributions of the avalanches and is predicted by:

$$\beta_{\text{pred}} = \frac{(\alpha - 1)}{(\tau - 1)}. \tag{A.3}$$

Comparing the fitted value from the empirical data ($\beta_{\text{fit}}$) and its estimation using $\alpha$ and $\tau$ exponents ($\beta_{\text{pred}}$), a new measure is derived to evaluate the *Deviation from Criticality Coefficient* (DCC), parameterised as $d_{CC}$:

$$d_{CC} = |\beta_{\text{pred}} - \beta_{\text{fit}}|, \tag{A.4}$$

where $\beta_{\text{pred}}$ and $\beta_{\text{fit}}$ are the predicted and fitted values of $\beta$ respectively. Consequently, a smaller DCC value indicates a more accurately fit power law distribution to the empirical data.

##### A.9 Branching ratio

The branching ratio is defined as the ratio of the number of units (neurons) active (spiking) at time step $t + 1$ to the number of active units (neurons) at time step $t$. Since a critical regime is naturally balanced and avoids runaway gains, the critical branching ratio is 1. Consequently, on average, network activity neither saturates nor dampens over time. Suppose that $N$ active neurons are detected in total and the number of active neurons in each time step $t$ is defined by $N(t)$. A fixed branching ratio of $m$, gives:

$$\langle N(t+1)|N(t)\rangle = mN(t) + h, \tag{A.5}$$

where $\langle|\rangle$ is the conditional expectation and $h$ is the mean rate of external drive. The activity decreases if $m < 1$, whereas it grows exponentially if $m > 1$, meaning that $m = 1$ separates these two regimens and represents a critical dynamic point. A precise prediction of $m$ helps to assess the risk that $N(t)$ will develop large and devastating avalanches of events such as epileptic seizures.

Under the circumstances when the full activity $N(t)$ is known, $m$ can be conventionally estimated using linear regression. Nevertheless, when using subsampling, when only a fraction of neurons in a neuronal network are sampled, this conventional method will be biased to some extent. The bias vanishes only if all units are sampled, because it is inherent to subsampling and cannot be overcome by obtaining longer recordings. Instead, inspired by the method introduced in [46] the subsampled activity $n(t)$ is utilized, where the fraction of recorded units to all cells is defined as a constant $\mu$. $n(t)$ here is a random variable whose expectation is proportional to the real $N(t)$ and $\langle n(t)|N(t)\rangle = \mu N(t) + \xi$, where $\mu$ and $\xi$ are constants. The bias value for the conventional linear estimator can now be calculated as:

$$m\left(\frac{\mu^2 \mathrm{var}(N(t))}{\mathrm{var}(n(t)) - 1}\right). \tag{A.6}$$

To overcome this subsampling bias, the method introduced by Wilting and Priesemann (2018) was utilized. Instead of directly using the biased regression of activity at time $t$ and $t + 1$, multiple linear regressions of activity between times $t$ and $t + k$ were performed with different time lags $k = 1, \ldots, k_{\max}$. Each of these $k$ values returns a regression coefficient $r_k$ with $r_1$ being equal to the result of a conventional estimator of $m$. With subsampling, all these regression slopes are biased by the same factor $b = \frac{\mu^2 \mathrm{var}(N(t))}{\mathrm{var}(n(t))}$. In these circumstances, instead of the exponential relation $r_k = m^k$ which is expected under full sampling, the equations generalizes to:

$$r_k = \frac{\mu^2 \mathrm{var}(N(t))}{\mathrm{var}(n(t))}.m^k = b.m^k. \tag{A.7}$$

Having multiple calculated $r_k$ values, both $b$ and $m$ are estimated, which are constant for all $k$.

Figure A5 compares the estimated branching ratio parameter from 6 different cultured cortical networks during a *Gameplay* and a *Rest* session.

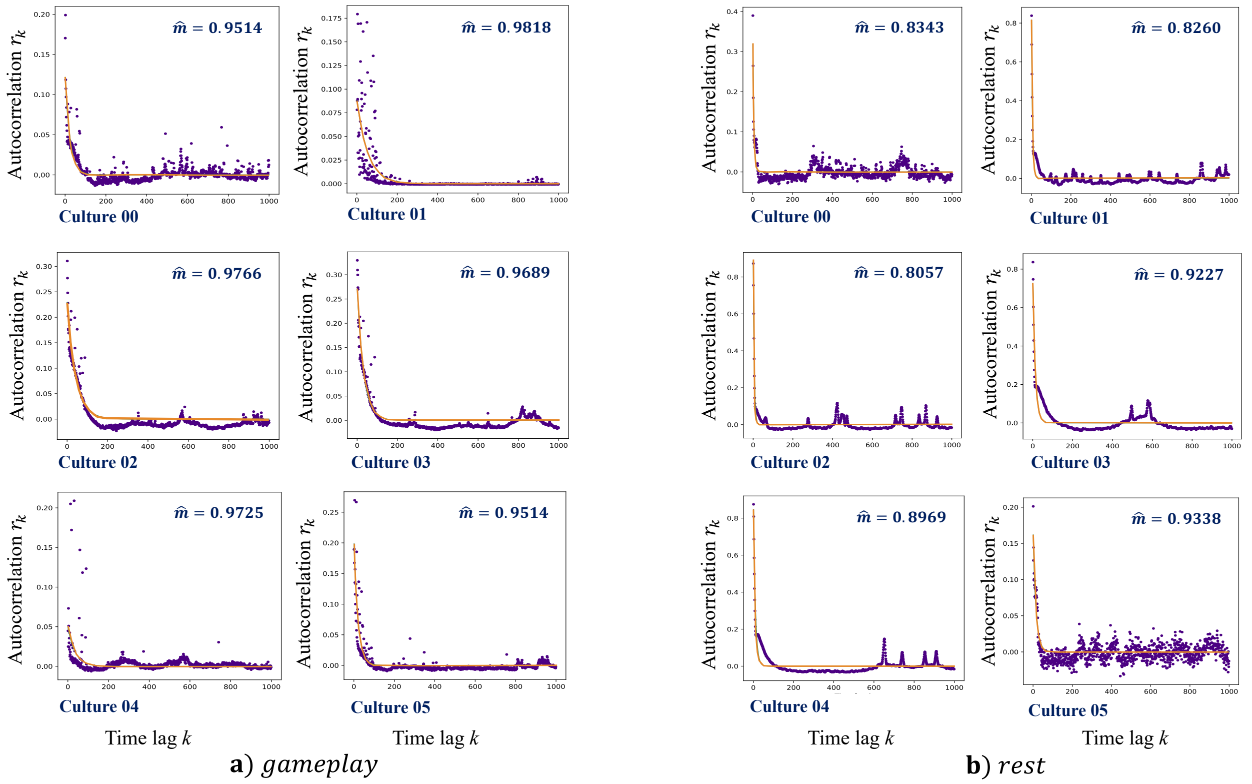


**Figure A5:** Estimated branching ratio parameters ($\hat{m}$) measured using the method introduced in [46] for 6 sample recorded cultures during a **a)** ***Gameplay*** session followed by a **b)** ***Rest*** session.

#### A.10 Scaling function

Another feature of critical dynamics is that avalanche shapes show fractal properties and all avalanche profiles of different sizes are scaled versions of the universal same shape. According to [66], the value of $\beta$ obtained from the exponent relation analysis can be used to calculate a scaling function for the avalanche shapes. For any given avalanche duration $D$, the average number of neurons firing at time $t$ (within $D$ seconds) is defined by $s(t, D)$. The following relations hold in this system:

$$
\begin{aligned}
s(t, D) &\propto D^{\gamma} F\left(\frac{t}{D}\right) \\
\langle S \rangle (D) &= \int_0^D s(t, D) dt,
\end{aligned}
\tag{A.8}
$$

where $F\left(\frac{t}{D}\right)$ is a universal function for all avalanches and $\gamma = \beta - 1$. Hence in this process, an initial $\beta$ is used to predict $\gamma$ and using this $\gamma$ and the first term in Equation A.8, $F\left(\frac{t}{D}\right)$ is obtained as $\langle \frac{s(t,D)}{D^{\gamma}} \rangle$. Here $\langle . \rangle$ denotes the average over all avalanches with duration $D$. A collection of $F\left(\frac{t}{D}\right)$ functions are extracted for various $D$ durations. The error for shape collapse is described as:

$$
\frac{\text{var}(F)}{\left(\max(F) - \min(F)\right)^2}. \tag{A.9}
$$

554 Repeating this process with various values for $\beta$, the exponent that produces the smallest shape collapse error in Equation A.9 is selected as the final scaling factor. In principle, we expect to obtain similar (if not the same) $\beta$ values from this analysis and the estimates in Section A.8. The NCC toolbox in MATLAB [47] was utilized to perform shape collapse on data. This shape collapse error is expected to be minimized under critical conditions. For shape collapse, avalanches with durations from 4 to 20 bins (200 to 1,000 ms) were considered. Across the time course of our recordings, there were not enough avalanches to conduct meaningful shape collapse analysis, beyond these cutoffs.

The schematic in Figure A6 summarizes the main attributes of a near-critical system compared to super/sub-critical states.

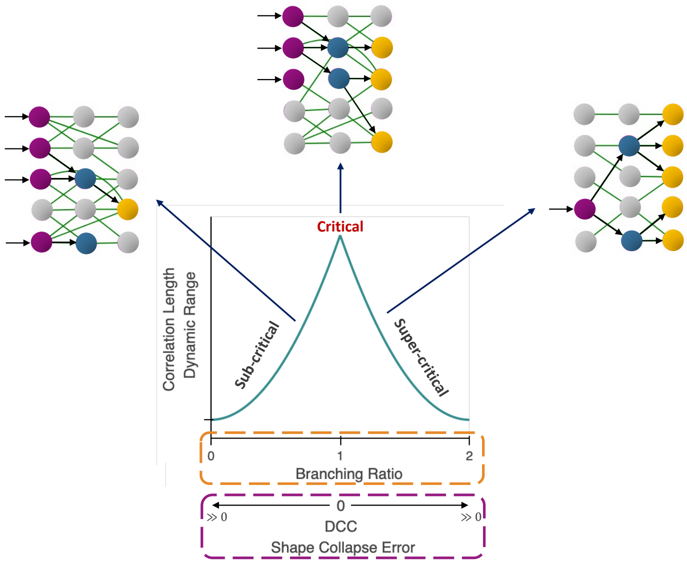


**Figure A6:** Key characteristics of a critical system compared with *sub-critical* and *super-critical* dynamics. If tuned near criticality, the branching ratio tends to 1 whereas the DCC and shape collapse error are close to 0.
